# Unfolding admixed ancestry and genomic diversity in zoo giraffes

**DOI:** 10.1101/2025.03.31.645993

**Authors:** Mischa F. Fraser, Rasmus Heller, Laura D. Bertola, Christina Hvilsom, Renzo F. Balboa, Xiaodong Liu, Anna Brüniche-Olsen

## Abstract

A giraffe nicknamed Marius attracted international media attention and world-wide notoriety when euthanized by Copenhagen Zoo in 2014, deemed surplus to the European breeding program. This incident highlights the challenges of managing zoo populations to best contribute to species conservation amid limited resources. Here, we investigated the degree of admixture, genetic diversity and inbreeding in *ex situ* giraffes using whole genome sequencing data from Marius and 12 other zoo giraffes, leveraging a reference data set from 71 wild giraffes. We found that most zoo giraffes show admixed ancestries. Marius, and zoo individuals assigned to Reticulated giraffes, exhibits mosaic ancestry with 85% Reticulated and 15% Nubian origin. We further detected recent inbreeding in some of the zoo individuals, consistent with their studbook data. Runs of homozygosity revealed that Marius and three other zoo giraffes were more inbred than their wild counterparts, although their admixed ancestry compensates for the reduction in genetic diversity caused by inbreeding. Our findings highlight the dual challenges of admixture and inbreeding in *ex situ* populations of giraffe, emphasizing the need for balanced genetic management to mitigate inbreeding and outbreeding degression while preserving diversity reflective of wild populations. This study also reinforces the importance of studbooks and genomic tools in guiding ex situ conservation strategies.

## Introduction

Zoos function as *ex situ* reservoirs that can be important assets in breeding programs and conservation of biological diversity. As part of maintaining healthy *ex situ* populations, many zoos participate in species breeding programs managed by regional zoo and aquarium associations. Allowing animals to breed is generally preferred overusing contraceptives from an animal welfare point of view (Asa et al. 2010). The captive breeding may lead to a surplus of animals—as zoos have limited space capacities—and these individuals are exchanged among zoos for breeding purposes to ensure genetic diversity is retained, or, in some cases, used for reintroductions and reinforcements of existing *in situ* populations, as demonstrated by species like the European bison (*Bison bonasus*) (Kleiman 1989) and the black-footed ferret (*Mustela nigripes*) (Dobson & Lyles 2000). However, reintroductions are challenging, costly and unfortunately not always successful (Schwartz et al. 2017).

Giraffes (*Giraffa camelopardalis*), classified as vulnerable by the IUCN, have experienced rapid population decline in the wild (Muller 2018), having more than halved since the 1980s with an estimated 70,000 individuals remaining. Giraffes are relatively abundant in zoos—close to 2,000 as of July 2024 (Species360 Zoological Information Management System (ZIMS), figure 1a)—due to their role in conservation education, awareness campaigns, charismatic appeal and relatively easy husbandry (Bertelsen 2014; Muller 2018; Foundation 2022). As a flagship species, giraffes generate significant public interest in conservation. This was evident in the global controversy surrounding the 2014 euthanasia of a 2-year-old giraffe—nicknamed Marius (GAN: MIG12-30090327)—at the Copenhagen Zoo, Denmark (figure 1b) (Clauss et al. 2025). Despite extensive media coverage and an online petition with 27,000 signatures to save him, Marius was euthanized, highlighting the potential conflicts between science, ethics and public opinion inherent in zoo breeding programs. For instance, every year zoos produce surplus individuals that are not included in future breeding, often due to their genetics already being (over-)represented in other individuals in the breeding programs (EAZA 2014; Crystal Allen 2018). Marius was part of this cohort of surplus animals (Clauss et al. 2025).

**Figure 1.**
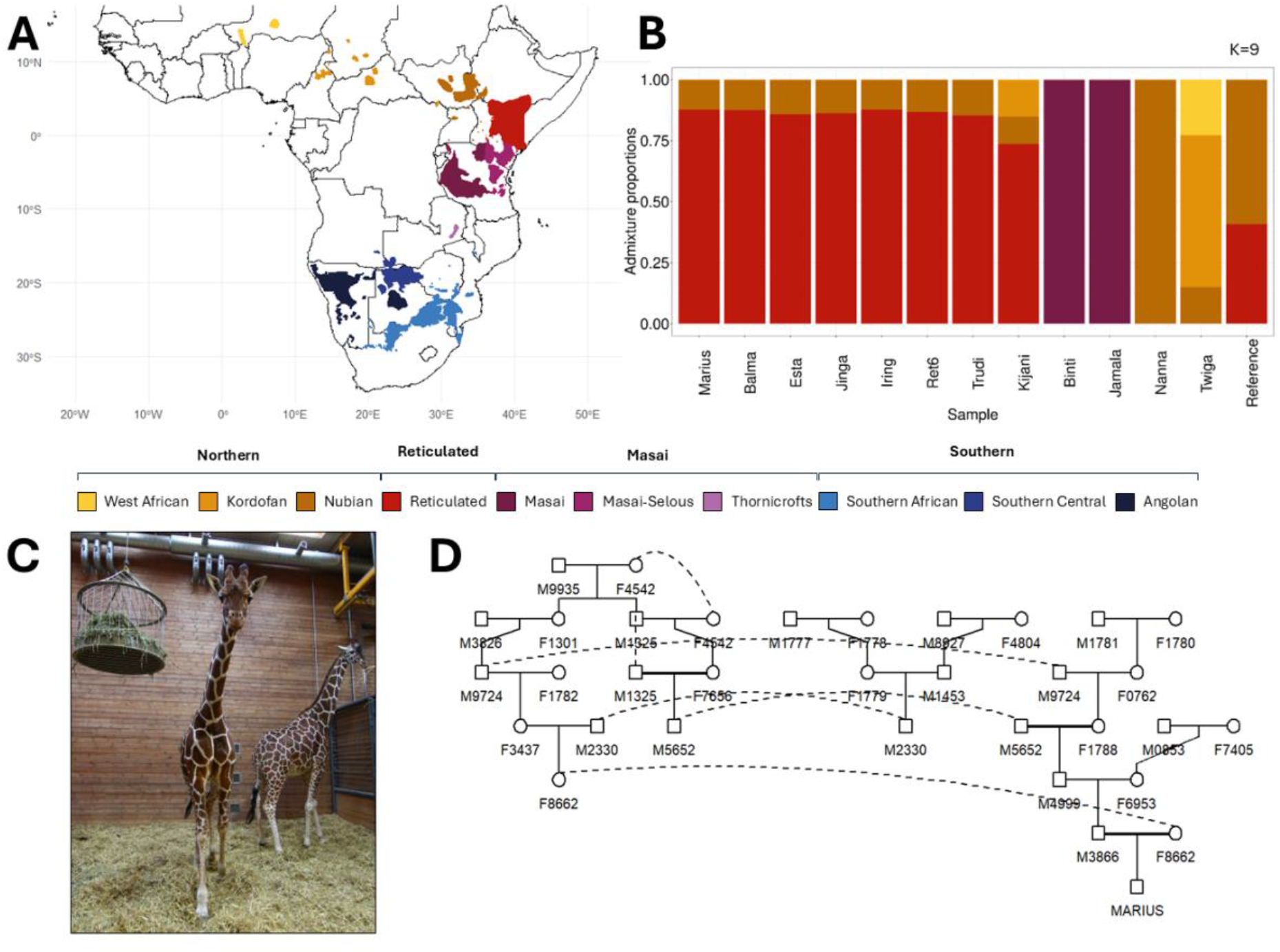
(A) Distributions of wild giraffe populations based on IUCN red list lineage ranges [5,18]. (B) Inferred ancestry proportions for zoo giraffes based on a supervised ADMIXTURE analysis in which nine giraffe lineages in the wild are used as the reference panel. (C) A photo of Marius on the left. (D) Pedigree of Marius based on the studbook information.

Molecular information is increasingly used in the management of *ex situ* species to prevent inbreeding, mitigate genomic load, and ensure that genomic lineages are maintained, and lineage specific traits are conserved (Frandsen et al. 2020; Jensen et al. 2020; Putnam et al. 2023; Speak et al. 2024). A series of recent studies have highlighted very strong population structures within giraffes (Winter et al. 2019; Coimbra et al. 2021b; Coimbra et al. 2022; Coimbra et al. 2023; Bertola et al. 2024; Prochotta et al. 2024), leading to the recognition of several distinct giraffe lineages, considered as separate species by some (figure 1a), although the exact number is debated (Winter et al. 2019; Coimbra et al. 2021b; Bertola et al. 2024). Among the lineages large variation in levels of genetic diversity and population sizes is observed, particularly giraffes in the southern part of the species range show low genetic diversity and high levels of inbreeding, making them relevant for conservation efforts (Bertola et al. 2024). Therefore, giraffe gene pools, including those contained in *ex situ* programmes such as zoos, should be carefully monitored and managed, as they potentially represent a valuable reservoir of genetic diversity. However, to our knowledge no studies has yet investigated the genetic composition of zoo giraffes. Here, we present analyses based on whole genome sequencing of Marius along with 12 additional zoo giraffes managed under the European Association of Zoos and Aquaria (EAZA) *Ex situ* Programme (EEP). We examined their genetic ancestry, genetic diversity and runs of homozygosity (ROHs), and compared these to an extensive dataset representing all major lineages of wild giraffes.

## Materials and methods

### Samples, studbook information and DNA data

We used newly generated DNA sequence data from two EEP individuals including Marius, together with SRAs from 11 EEP giraffes including one from Agaba et al. (2016), one from Farré et al. (2019), eight from Coimbra et al. (2021b), and one from Liu et al. (2021) (figure S2). Additionally, we included DNA sequence data from 71 unrelated wild giraffes belonging to nine discrete populations and two okapis from Bertola et al. (2024). Data from samples were processed following Bertola et al. (2024). In brief, we mapped the sequence data to the reference genome of giraffe (GenBank: GCA_017591445.1) using the PALEOMIX pipeline (Schubert et al. 2014). We next used BCFtools (Danecek et al. 2021) to perform variant calling. The resulting polymorphic dataset only contains bi-allelic SNPs from the genomic regions passing the site filters generated in Bertola et al. (2024), in which low complexity sequences and paralogous loci were removed. Based on this SNP dataset, we estimated relatedness for all 13 captive giraffes using R0, R1 and KING-robust estimator (Waples et al. 2019). To assess if the studbook information matched the genomic results, we further obtained the studbook ancestry information for all 13 captive giraffes.

### Genetic origins and admixture

To assess the genetic ancestry of the captive individuals we conducted a supervised ADMIXTURE analysis to estimate their admixture proportions using FastNGSadmix (Jørsboe et al. 2017) on two levels. At the species level, we used the four major lineages as the reference panel including Reticulated, Masai, Southern and Northern giraffes. At the population level, we used the nine populations identified in Bertola et al. (2024) as the reference panel, representing a comprehensive dataset of giraffe populations *in situ*. To further validate the admixture patterns observed in some captive samples, we calculated *D*-statistics using the function qpDstat from ADMIXTOOLS2 (Maier et al. 2023). We used the Reticulated population as H1, captive sample to test as H2, the Nubian population as H3 and the Okapi as an outgroup to specifically test presence of admixture between the Nubian populations and captive individuals assigned to ‘Reticulated’.

### Heterozygosity and runs of homozygosity

We assessed genetic diversity of the captive giraffes based on levels of individual genome-wide heterozygosity and runs of homozygosity (ROH) and compared these results with their wild counterparts. Individual heterozygosity was estimated based on called genotypes as the proportion of heterozygous sites compared to all sites (variable and non-variable) passing the site filters. We next called ROH using the “--homozyg” function in PLINK v1.9 (Purcell et al. 2007) with default settings, except for allowing 3 heterozygous SNPs (--homozyg-window-het 3) and 20 missing calls (--homozyg-window-missing 20) within the scanning window. To prepare the input, we applied additional filters to the SNP dataset. These filters include masking heterozygous calls of distorted allelic balance (AB<0.25 or AB>0.75) and removal of SNPs with a maximum minor allele frequency of 0.05, a minimum data missingness of 5% and a minimum ratio of observed heterozygotes of 50% within each subspecies. To investigate how recent inbreeding has impacted the diversity of these individual giraffe samples, we also estimated individual heterozygosity excluding the respective ROH regions.

## Results

### Ancestry and admixture

A supervised ADMIXTURE analysis on the population level demonstrated varying degrees of genetic admixture in zoo giraffes (figure 1b, figure S1). Only three individuals displayed unadmixed ancestry with respect to the wild populations: Binti (Masai), Jamala (Masai) and Nanna (Nubian). In contrast, Kijani and Twiga displayed complex patterns of admixture involving three lineages; Kijani showed admixture between the Reticulated (∼75%), Nubian (∼10%) and Kordofan (∼15%) lineages, while Twiga shows admixture between the Kordofan (∼15%), Nubian (∼65%) and West African (∼20%) lineages. The individual representing the NCBI reference genome assembly of a Rothschild’s giraffe (classified within the Nubian lineage) exhibited admixture between the Nubian (∼60%) and Reticulated (∼40%) giraffe lineages. Notably, several EAZA individuals designated as Reticulated giraffes in their studbook, including Marius (figure 1c,d), displayed similar patterns of admixed ancestries with approximately 85% Reticulated and 15% Nubian (∼15%) lineage contributions. These similarities in admixture proportions very likely result from shared ancestors in their pedigrees, as supported by studbook pedigrees (figure S2), and high levels of relatedness among the Reticulated Zoo giraffes, in which five out of eight individuals are involved in five relative pairs of first degree or second degree (figure S3). The admixture patterns observed in the zoo individuals designated as Reticulated giraffes were further corroborated by *D*-statistics, which identified significant signals of introgression between these zoo giraffes and the Nubian lineage (figure S4).

### Genomic diversity and inbreeding

The zoo individuals with predominant ancestry of the Reticulated lineage show higher genome-wide heterozygosity than other *ex situ* giraffes, mirroring the situation in the wild where Reticulated giraffes have higher genetic diversity than other lineages (figure 2a) (Bertola et al. 2024). In general, the zoo giraffes with admixed ancestries have higher heterozygosity compared with the average heterozygosity levels observed in their wild counterparts. Among the three non-admixed individuals, Binti and Jamala exhibit slightly higher heterozygosity than the Masai giraffe, whereas Nanna shows a slight decrease in heterozygosity compared to the Kordofan giraffe (figure 2a).

**Figure 2.**
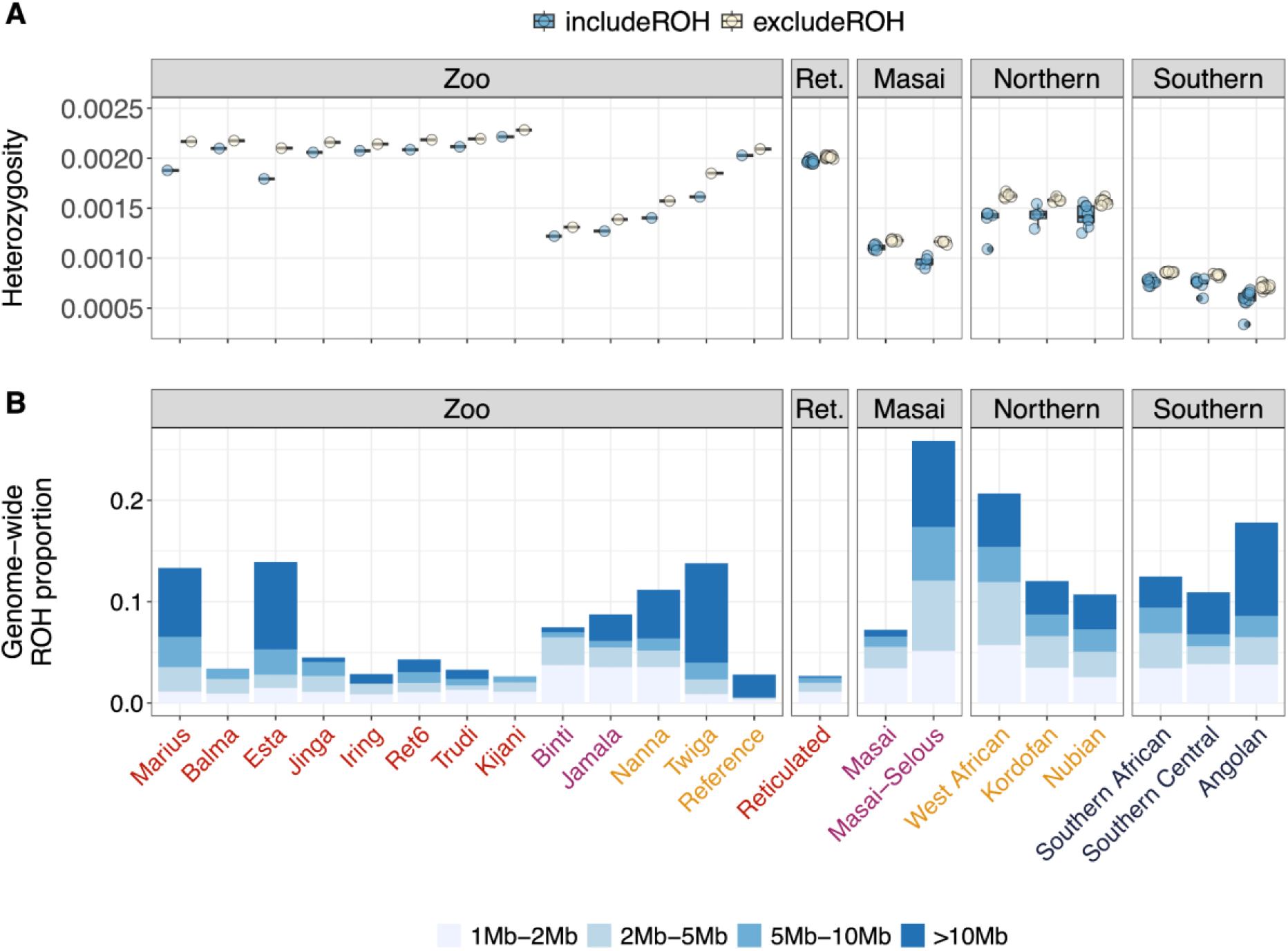
(A) Genome-wide heterozygosity including and excluding runs of homozygosity (ROH) for the zoo giraffe individuals and wild populations. (B) Genome-wide ROH proportions for zoo individuals and mean ROH in the wild populations. Proportions of differing ROH length intervals are shown as subdivisions within bars. The names of zoo giraffe individuals along the X-axis are colored according to the four major lineages that they are designated as, including Reticulated lineage, Masai lineage, Northern lineage and Southern lineage (see table S2).

We observed significant variation in ROH patterns among the zoo individuals (figure 2b). While individuals with Reticulated ancestry generally have few and short ROHs mirroring their wild counterparts, Marius and Esta show a large fraction of total ROH, especially long ROH (>10Mb) indicative of recent inbreeding (figure 2b, figure S5). In addition, Twiga, despite having multiple sources of ancestry, also exhibits excessive amounts of long ROH. These three individuals further show the most prominent differences between overall heterozygosity and heterozygosity excluding ROH, which indicates that their decreased diversity is mainly driven by recent inbreeding (figure 2a). The individual used to generate the reference genome has the smallest proportion of shorter (<10Mb) ROHs and mainly long (>10Mb) ROHs, consistent with very recent inbreeding (figure 2b) and a mosaic ancestry.

High genetic inbreeding in these zoo giraffes was further supported by their pedigree information derived from the studbooks, which presented several cases of mating among relatives, with individuals being used for breeding several times within families increasing autozygosity (figure S2). Reviewing Marius’s pedigree, both sire M9724 and dam F4542 appear twice (figure 1d). Furthermore, parents have bred with their offspring in several cases, e.g. Marius’s great-great-great-great grandparents (dam: F4542 and sire: M1325) and great-great-great grandparents (sire: M1325 and dam: F7656).

## Discussion

Our study represents the first genome-wide investigation of admixture, genetic diversity and inbreeding based on 13 zoo giraffe individuals by leveraging genomic data of *in situ* giraffe populations. Indeed, the increasing availability of giraffe genome data from populations across the species range (Coimbra et al. 2021a; Liu et al. 2021; Bertola et al. 2024; Prochotta et al. 2024) has the potential to genetically inform *ex situ* giraffe management. For such efforts to succeed, a comprehensive reference panel representing the genetic diversity in the wild is critical for accurately detecting ancestry and admixture of the *ex situ* populations (Hvilsom et al. 2013; Shringarpure & Xing 2014; Frandsen et al. 2020).

Genetically informed *ex situ* conservation faces similar basic challenges to *in situ* conservation: how to optimise the balance between inbreeding depression and outbreeding depression while still reflecting the diversity found in the wild (Frankham et al. 2011; Willi et al. 2022). The risk of inbreeding is often exacerbated in zoos, where relatively small population sizes make the matings between relatives hard to avoid (Boakes et al. 2007). Our study highlights this for some zoo giraffes, especially Marius and Esta representing the Reticulated giraffe. While it was beyond the scope of this study to investigate whether the observed levels of inbreeding leads to inbreeding depression, most of the zoo giraffes investigated here had higher inbreeding coefficients than their wild counterpart population (figure 2B). On the other hand, admixture among zoo giraffes has increased the heterozygosity of the zoo giraffes above what is found in their wild counterpart populations, except in Marius and Esta, where inbreeding is extensive enough to override the compensatory effect of admixture (figure 2A).

While admixture can mitigate the effects of inbreeding by introducing new genetic variation, it can also potentially lead to outbreeding depression (Frankham et al. 2011; Dussex et al. 2023). The extent of this risk is hard to assess theoretically. The rare occurrence of admixture in *in situ* giraffe populations despite their close geographic proximity (Bertola et al. 2024) is consistent with reproductive barriers between the major lineages of giraffes (Prochotta et al. 2024), but confinement to refugia with subsequent spatial expansion is an alternative explanation for the paucity of admixture *in situ*. To our knowledge, there are no indications of outbreeding depression in zoo giraffes, and recent research showed large amounts of historical admixture between the major giraffe lineages (Bertola et al. 2024). In summary, while this study cannot determine the functional implications of elevated inbreeding and admixture in zoo giraffes, it demonstrates the presence of both processes and the need to account for potential risks of inbreeding and outbreeding depression in future *ex situ* management strategies. Specifically, we find that all examined zoo giraffes designated as the Reticulated giraffe were admixed, and individuals like Marius and Esta may simultaneously face compounded risks of both inbreeding and outbreeding depression. These findings warrant further, larger-scale genetic investigations of the *ex situ* giraffe population with an emphasis on individuals representing Reticulated giraffes.

Our findings also highlight the challenges relying on misaligned metadata in public repositories. DNA from zoo animals is often used for genetic studies, including generating reference genome assemblies. However, many of the zoo giraffes examined here were not purely representative of the wild lineages they were assigned to. For example, the current highest-quality giraffe reference genome (GenBank assembly: GCA_017591445.1), designated as a Rothschild’s (Nubian) giraffe (Liu et al. 2021), is heavily admixed with Reticulated giraffe (figure 1b). Such misassignment can lead to errors in taxonomic (Keck et al. 2023) and genomic studies (Gopalakrishnan et al. 2017; Thorburn et al. 2023) if not accounted for.

Our results illustrate how extensive knowledge of the genetic structure *in situ* populations can facilitate the application of genomics for determining origin and admixture proportions of *ex situ* individuals. Additionally, genomics can provide critical insights when studbook records are unavailable, as demonstrated by the individual from Guangzhou Zoo used to create the giraffe reference genome (Liu et al. 2021). Importantly, our study highlights the balance between inbreeding- and outbreeding depression as a key challenge in the management of zoo giraffes. While our findings provide valuable insights, it is important to acknowledge that the scope of this study is limited by sample size. Nonetheless, our study is based largely on random sampling across multiple zoos, suggesting that these results are likely representative of broader trends in ex situ giraffe populations. These findings highlight the necessity of further large-scale genetic investigations into the ex situ giraffe population and provide a baseline for future studies. Finally, our results show that integrating large-scale genetic assessments into management plans and breeding programs can provide important baseline data for *in situ* populations.

## Data and code accessibility

DNA sequence data of Marius (GAN: MIG12-30090327) and Nanna (GAN: MIG12-29806330) have been uploaded to the following NCBI repository: PRJNA1243753. Already published DNA sequence data can be found in these repositories: PRJEB66216, PRJNA635165, SRR3218456, SRS3530193, GCA_017591445.1. All code has been uploaded to GitHub (https://github.com/popgenDK/seqAfrica_Marius).

## Acknowledgements

We thank Copenhagen Zoo and Aalborg Zoo for providing blood samples from two giraffes, Marius and Nanna from which much of this work revolved. We thank Sven Winter for providing studbook information for zoo giraffe individuals from previous publications. We thank Frank Rønsholst for providing official photographs. This work was supported by the Department of Atomic Energy, Government of India, Project Identification No. RTI 4006.

