## Supplemental Figures and Tables for "Unfolding admixed ancestry and genomic diversity in zoo giraffes"

### Supplemental Information

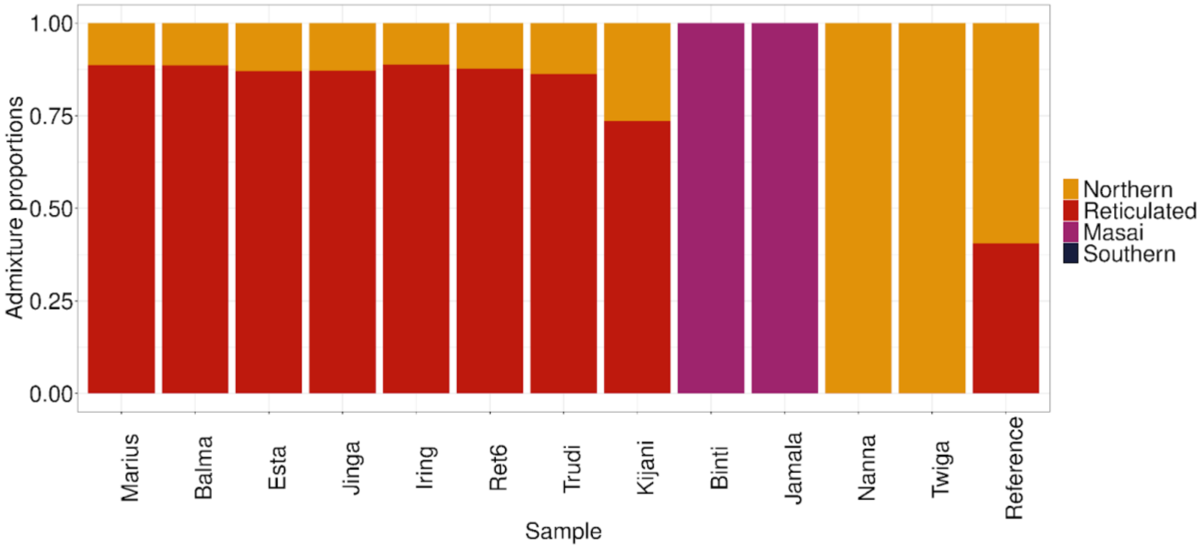


**Supplementary Figure S1.** Ancestry proportions for the 13 zoo giraffes based on a supervised ADMIXTURE analysis, where the four major lineages of giraffes (in the legend) are used as the reference panel.

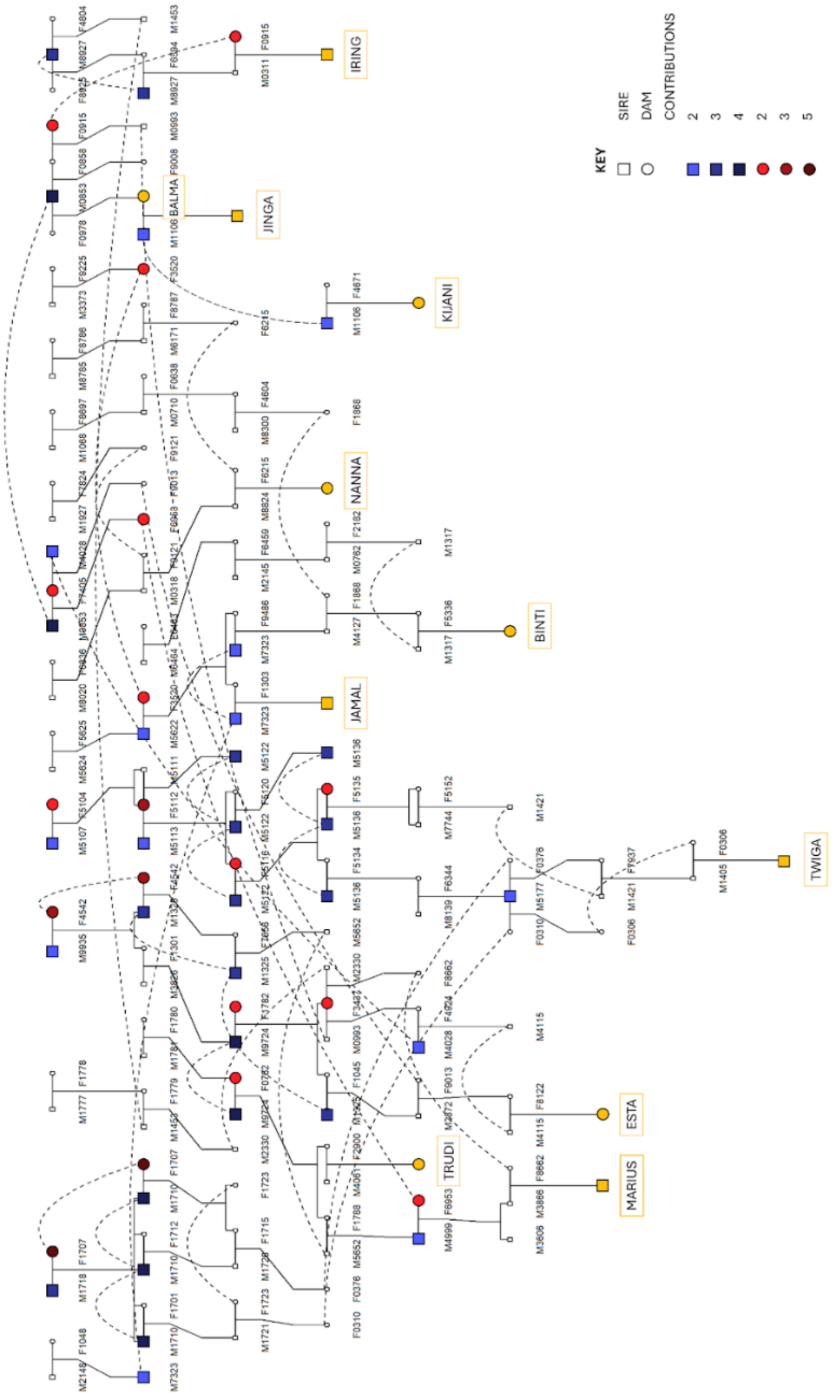


**Supplementary Figure S2**. Kinship2 pedigrees of zoo giraffes in which Studbook information is available.

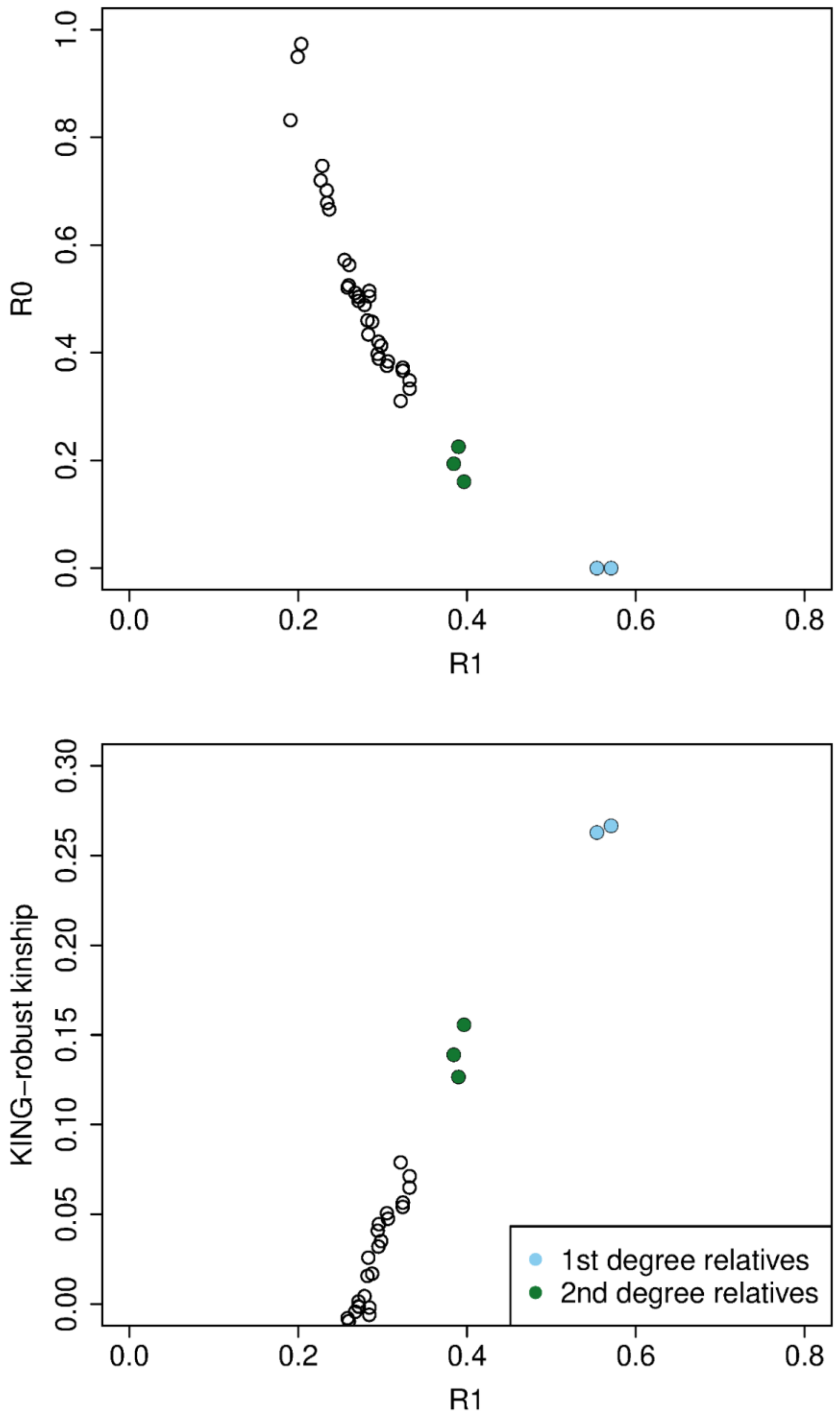


**Supplementary Figure S3**. Relatedness estimation for the 13 zoo giraffes following Waples et al. [36]. We identified two pairs of 1st degree relatives: Balma-Jinja and Balma- Ret6, and three pairs of 2nd degree relatives: Balma- Trudi, Iring- Trudi and Ret6- Trudi.

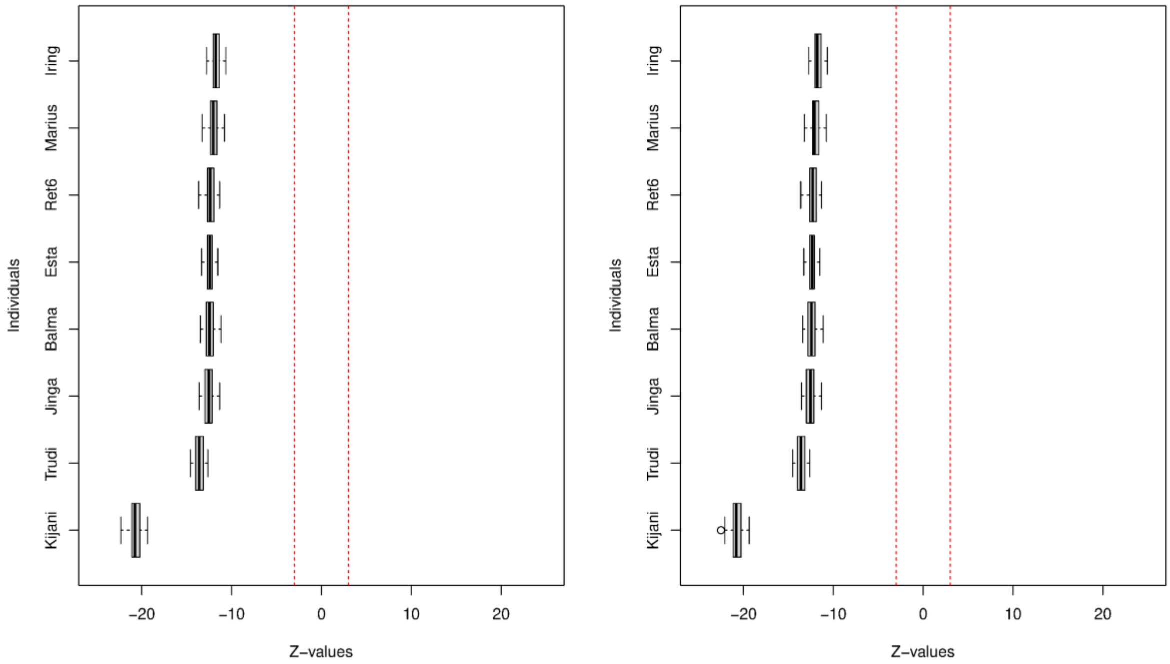


**Supplementary Figure S4**. Summary of the *D*-statistics, in which we included the Reticulated lineage as H1, the zoo individuals to test as H2 (on the Y axis), the Nubian lineage as H3 and the Okapi as outgroup. This setup allowed us to specifically test for admixture between the zoo giraffes and Nubian lineage. The red dotted lines in the figure indicate an absolute Z-value of 3.

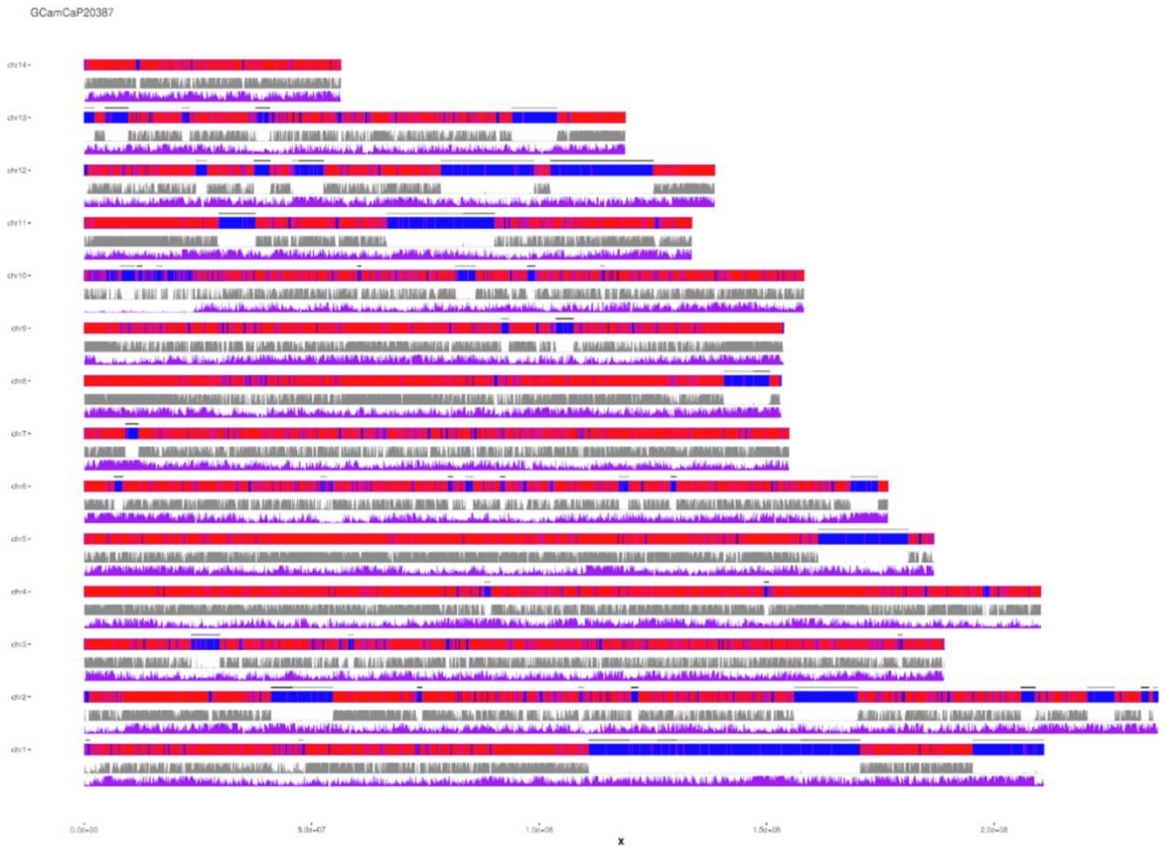


**Supplementary Figure S5**. Genotype calls, proportions of heterozygous sites and SNP density for ROH validation in Marius. For each of the 14 chromosomes homozygous sites (blue) and heterozygous sites (red)r are depicted. Identified ROHs are marked by black horizontal bars. The gray line below homozygous/heterozygous calls shows proportions of heterozygous calls in a window of 100 kb. The purple line below shows the number of SNPs along the 100 kb window.

**Table S1**. Giraffe holdings from Species 360: Zoological Information Management System (ZIMS; accessed July 2024). Two lineages discussed in Bertola et al. (2024) (West African and Thornicrofts) are not listed in ZIMS; Nubian is called Rothschild's in ZIMS.

| | **Major lineage** | **Male** | **Female** | **Unknown** | **Total** |
|---|---|---|---|---|---|
| **Generic** | | 235 | 341 | 1 | **577** |
| **West African** | Northern | - | - | - | - |
| **Kordofan** | Northern | 45 | 44 | 1 | **90** |
| **Nubian** | Northern | 221 | 304 | 0 | **525** |
| **Reticulated** | Reticulated | 205 | 338 | 17 | **560** |
| **Masai** | Eastern | 62 | 78 | 0 | **140** |
| **Thornicroft** | Eastern | - | - | - | - |
| **Transvaal** | Southern | 26 | 48 | 0 | **74** |
| **Angola** | Southern | 9 | 6 | 0 | **15** |
| **Total** | | **803** | **1159** | **19** | **1981** |

**Table S2.** Sample information of both zoo and wild giraffes by GAN (global accession number), lineage, paper ID and reference.

| **Name** | **GAN** | **Assigned giraffe lineage (major lineage)** | **PaperID** | **Reference** |
|---|---|---|---|---|
| Marius | MIG12-30090327 | *G.c.reticulata* (Reticulated) | Marius | this study |
| Balma | 5669004 | *G.c.reticulata* (Reticulated) | RET1 | [15] |
| Esta | 24937489 | *G.c.reticulata* (Reticulated) | TRot2 | [15] |
| Jinga | 26966871 | *G.c.reticulata* (Reticulated) | RET4 | [15] |
| Iring | 5669002 | *G.c.reticulata* (Reticulated) | RET5 | [15] |
| Ret6 | 26962683 | *G.c.reticulata* (Reticulated) | RET6 | [15] |
| Trudi | MIG12-29316778 | *G.c.reticulata* (Reticulated) | TRot1 | [15] |
| Kijani | 26966872 | *G.c.reticulata* (Reticulated) | RET3 | [15] |
| Binti | MIG12-29882484 | *G.c.tippelskirchi* (Masai) | NZOO | [20] |
| Jamala | CWF10-00169 | *G.c.tippelskirchi* (Masai) | R1865 | [21] |
| Nanna | MIG12-29806330 | *G.c. rothschildi* (Northern) | Nanna | this study |
| Twiga | SVT18-03769 | *G.c.antiquorum* (Northern) | PLA01 | [15] |
| Reference | - | *G.c. rothschildi* (Northern) | rggzc | [22] |
